# Complement factor C4a does not activate protease activated receptor 1 (PAR1) or PAR4 on human platelets

**DOI:** 10.1101/2020.06.01.127662

**Authors:** Xu Han, Maria de la Fuente, Marvin T. Nieman

## Abstract

**Background:** Protease activated receptor 1 (PAR1) and PAR4 are key thrombin signal mediators for human platelet activation and aggregation in response to vascular injury. They are primarily activated by thrombin cleavage of the N-terminus to expose a tethered ligand. In addition to the canonical activation by thrombin, a growing panel of proteases can also elicit PAR1- or PAR4-mediate signal transduction. Recently, complement factor C4a was reported as the first endogenous agonist for both PAR1 and PAR4. Further, it is the first endogenous non-tethered ligand that activates PAR1 and PAR4. These studies were conducted with human microvascular cells; the impact of C4a on platelet PARs is unknown.

**Objectives:** The goal of this study was to interrogate PAR1 and PAR4 activation by C4a on human platelets.

**Methods:** Platelet rich plasma (PRP) were isolated from healthy donors. PRP was stimulated with C4a and the platelet aggregation was measured. HEK293 Flp-In T-rex cells were used to further test if C4a stimulation can initiate PAR1- or PAR4-mediated Gα_q_ signaling, which was measured by intracellular calcium mobilization.

**Results:** C4a failed to elicit platelet aggregation via PAR1- or PAR4-mediated manner. In addition, no PAR1- or PAR4-mediated calcium mobilization was observed upon C4a stimulation on HEK293 cells.

**Conclusions:** Complement factor C4a does not activate PAR1 or PAR4 on human platelets. These data show that PAR1 and PAR4 activation by C4a on microvascular cells likely requires a cofactor, which re-enforces the concept of cell-type specific regulation of protease signaling.

**Essentials:**

- C4a is an agonist for both PAR1 and PAR4 on human microvascular cells.
- We sought to determine if C4a activates human platelets through PAR1 or PAR4.
- C4a does not activate PAR1 and PAR4 on human platelets.
- This re-enforces the concept of cell-type specific regulation of PAR signaling.

## 1. Introduction

Protease activated receptors are the primary means by which proteases mediate intracellular signaling. The prototypical receptor, PAR1, was identified 30 years ago as the thrombin receptor.(1) Subsequently, three other family members were identified (PAR2, PAR3, and PAR4) by homology screening of cDNA libraries.(2) PARs are G-protein coupled receptors (GPCRs) that have a unique activation mechanism in which the N-terminus is cleaved by the activating protease to generate a tethered ligand. The newly exposed ligand binds intramolecularly to the receptor, inducing a conformational change that initiates signal transduction.(3) In addition to canonical thrombin-mediated signaling, there is a growing panel of identified proteases that also have the ability to activate PARs.(4) PAR1 can be cleaved by activated protein C (APC), matrix metalloproteinase 1 (MMP-1), MMP-2 and MMP-13 at non-canonical sites to generate distinct tethered ligands.(2, 3) These ligands initiate diverse signaling pathways depending on the cell type and cellular cofactors.(4) In addition to thrombin, PAR4 can be activated by trypsin, plasmin, cathepsin G, factor Xa, however, these all cleave at the canonical thrombin site and results the same downstream signaling.(3, 5, 6)

Human platelets express PAR1 and PAR4.(3) This dual receptor system works in concert to provide the sensitivity and duration of signaling required for hemostasis.(7) PAR1 is highly sensitive to thrombin, and results in a rapid, transient signal. In contrast, PAR4 responds to higher concentrations of thrombin and leads to a prolonged signaling response. Thrombin activation of PARs initiates multiple signaling cascades in human platelets by directly coupling to Gα_q_ and Gα_12/13_.(8) Gα_q_ stimulates the formation of inositol triphosphate (IP3) and diacylglycerol (DAG), leading to intracellular calcium mobilization and protein kinase C (PKC) activation.(3) Gα_12/13_ mediates Rho guanine nucleotide exchange factors and RhoA activation, which controls platelet shape change, spreading, and thrombus stability.(3) In platelets, PAR1 and PAR4 signal indirectly through Gα_i_, via secondary stimulation of the Gαi-coupled P2Y12 receptor due to released ADP.(3) However, Voss and colleagues used a pharmacological approach to show that PAR1 signals directly through Gαi in platelets.(9) In sum, PAR1 and PAR4 mediate several important signaling events in platelets.

The complement and coagulation systems communicate with one another.(10, 11) Platelets can lead to the activation of complement factors C3 and C5. Further, C4 activation can occur on platelets.(10, 11) Recently, Wang *et al.* have demonstrated that C4a is a novel soluble, untethered agonist for both PAR1 and PAR4 on human microvascular cells (HMEC).(12) Given the importance of PAR1 and PAR4 for platelet physiology and the crosstalk between the complement system and platelets, we explored the potential of C4a as an agonist for human platelets.

## 2 Methods

### 2.1 Reagents

The V5-tag antibody conjugated with FITC (catalog # R963-25) was from Invitrogen (Carlsbad, CA). Trypsin (catalog # V511A) was from Promega (Fitchburg, WI). The PAR1 activation peptide, SFLLRN-NH_2_ and TFLLRN-NH_2_ were from Tocris Bioscience (Minneapolis, MN). PAR4 activation peptide, AYPGKF-NH_2_, was from GenScript (Pisataway, NJ). C4a (Catalog # A106, Lot#16) was from Complement Technology (Tyler, TX). Vorapaxar was from Axon Medchem (catalog # 1755, batch 2). All other reagents were from Thermo Fisher Scientific (Pittsburgh, PA) except where noted.

### 2.2 Platelet isolation and aggregation

With approval by the Case Western Reserve University IRB, human whole blood was obtained by venipuncture from healthy donors after obtaining informed consent. Whole blood was collected into sodium citrate (3.2% buffered sodium citrate). The blood was centrifuged at 200 x *g* for 15 min to obtain PRP. Platelet concentrations were quantified using a Coulter Counter. PRP was stimulated with AYPGKF-NH_2_ (250 μM), SFLLRN-NH_2_ (30 μM), or C4a (3.2 μM). Platelet aggregation was measured under constant stirring (1200 rpm) with a Chrono-log Model 700 aggregometer using Aggrolink8 version 1.3.98.

### 2.3 Cell culture

As reported previously, HEK293 Flp-In T-rex cells were maintained in Dulbecco’s modified Eagle’s medium (Invitrogen) containing 5% fetal bovine serum (FBS) at 37 °C with 5% CO_2_. Human PAR4 containing an N-terminal V5-epitope were stably expressed using the HEK293 Flp-In T-Rex cells according to the manufacturer’s protocol (Invitrogen). The concentration of tetracycline was titrated (700 ng/mL) to yield PAR4 expression at ~175,000 receptor on the cell surface at 40 hr as determined by quantitative flow cytometry as described below.(13) Some cells were processed for flow cytometry and the remaining cells were processed for Ca^2+^ mobilization.

### 2.4 Flow Cytometry

Following induction, PAR4 expressing cells were harvested in Versene solution and collected by centrifugation. The cells were then washed 3 times with HEPES-Tyrode’s Buffer, pH 7.4 (10 mM HEPES, 12 mM NaHCO_3_, 130 mM NaCl, 5 mM D-glucose, 5 mM KCl, 0.4 mM NaHPO_4_, 1 mM MgCl_2_). To assess PAR4 expression prior to each signaling experiment, cells were incubated with FITC conjugated anti-V5 antibody for 30 minutes. Cells were analyzed using a BD LSRFortessa™ cell analyzer (BD Biosciences, Mississauga, ON, Canada). PAR4 expression was quantified using Quantum Simply Cellular beads from Bangs Laboratories, Inc. (Fishers, IN) to generate a standard curve of antibody binding sites as previously described.(13)

### 2.5 Calcium mobilization

As reported previously, parental HEK293 Flp-In or PAR4 expressing cells were incubated with 0.5 μM Fura-2 with or without 100 nM vorapaxar for 1 hr. The cells were then washed 3 times in HEPES-Tyrode’s buffer with 200 μM CaCl_2_ and diluted to 2.5 × 10^5^ cell/mL. The intracellular calcium mobilization was recorded using a fluorometer from Photon Technology International Inc for 420 s total. The first 50 s was set as the background, the response to agonist was measured for 250 s, and the maximum and minimum fluorescence was measured for 60 s each by adding 0.1% Triton X100 and 8.8 mM EDTA sequentially.(13) Fluorescence values were converted to intracellular free Ca^2+^ concentration with the equations described by Grynkiewicz et al.(13, 14)

## 3. Results and Discussion

Wang et al used a cell-based β-arrestin reporter assay to demonstrate that C4a dose-dependently activated both PAR1 and PAR4 in HMEC-1.(12) The EC_50_ of C4a activation of PAR1 and PAR4 was 0.8 μM and 0.6 μM, respectively.(12) Since PAR1 and PAR4 have primary roles in platelet activation, we investigated the potential of C4a to serve as an agonist for human platelets. Platelet rich plasma (PRP) was isolated from whole blood of two independent donors on different days for aggregation studies. In both cases, platelets responded to 30 μM PAR1-AP (SFLLRN) or 250 μM PAR4-AP (AYPGKF) (**Figure 1**). In contrast, platelets did not respond to 3.2 μM C4a (**Figure 1**). We extended the recording time to 16 minutes and still no aggregation was triggered by 3.2 μM C4a (**Figure 1B**). The EC_50_ values of C4a reported by Wang et al is 0.8 μM for PAR1 and 0.6 μM for PAR4, which is 4 or 5.3-fold lower to the dose we treated human platelets, respectively.(12) Further, the plasma concentration of C4a varies from 68.91 ±33.23 ng/ml (7.97 nM) measured by a cytometric bead assay to 2398 ng/ml (277 nM) using an ELISA.(15, 16) Nonetheless, in our experiments, platelets do not respond to C4a at a concentration 12 – 400 fold higher than is found in plasma.

**Figure 1:**
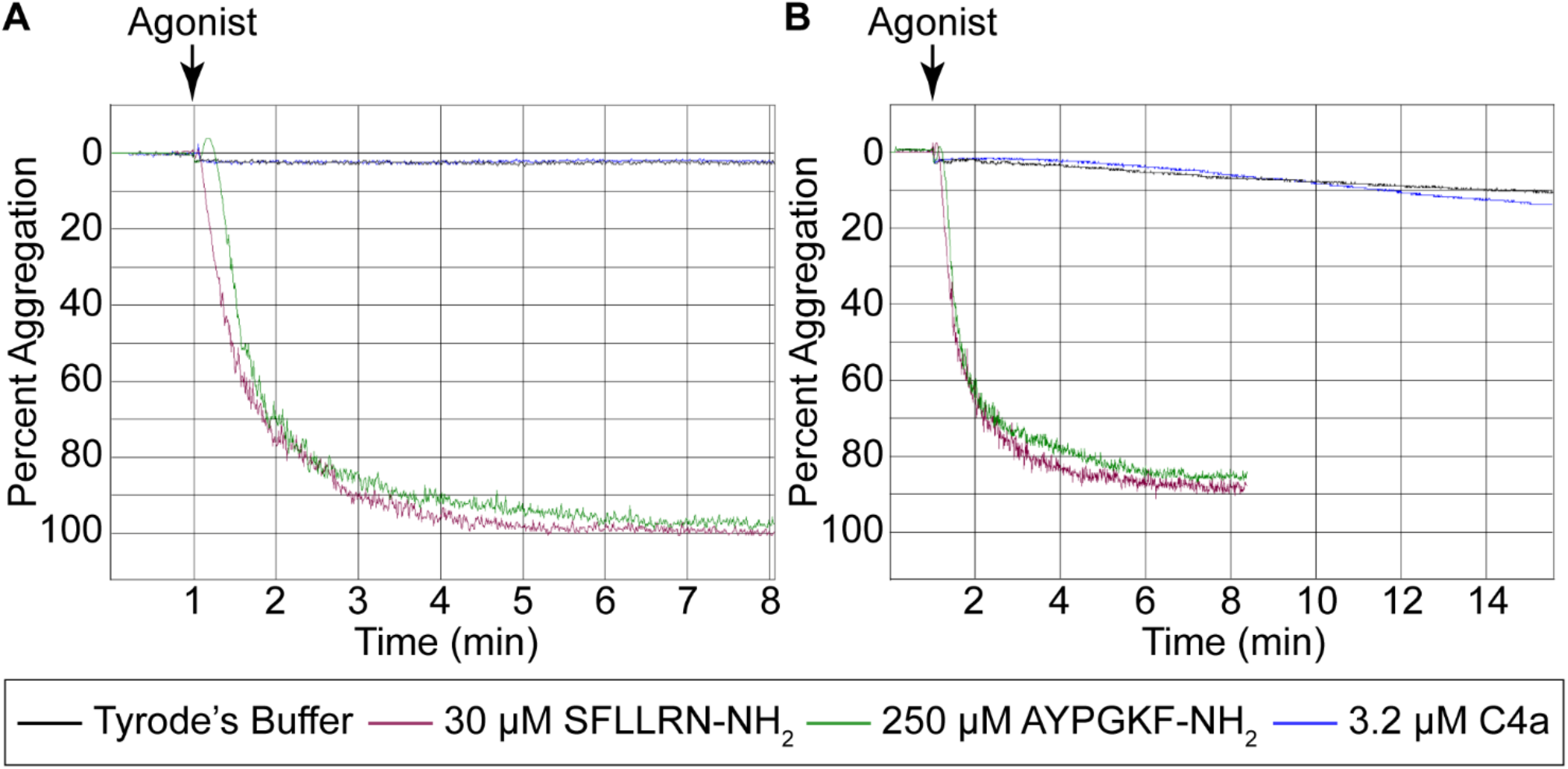
C4a failed to trigger platelet aggregation. Human platelet rich plasma (PRP) was isolated from independent healthy donors (A and B). Following stimulation with Tyrode’s buffer (Black trace), SFLLRN (30 μM) (Red trace), AYPGKF (250 μM) (Green trace), or C4a (3.2 μM) (Blue trace), percent aggregation was determined. In Panel B the platelets were monitored for an addition 8 min following stimulation.

We next tested the ability of C4a to activate calcium signaling via PAR1 in HEK293 Flp-in T-rex cells, which express PAR1 endogenously. Thrombin or the PAR1-specific activation peptide, TFLLRN-NH_2_, induced transient intracellular Ca^2+^ mobilization (**Figure 2A and B**, red traces). In both cases, Ca^2+^ signaling was completely abolished when the cells were preincubated for 1 hour with 100 nM of PAR1 antagonist, vorapaxar (**Figure 2A and B**, orange traces). As expected, there was no calcium flux induced by the stimulation of 250 μM of PAR4-activation peptide (AYPGKF-NH_2_) since HEK293 Flp-in cells do not express PAR4 endogenously (**Figure 2B**). We next used this experimental design to specifically test PAR1 response to C4a. Similar to our platelet aggregation experiments, C4a did not activate PAR1-mediated Gα_q_ signaling at 3.2 μM (**Figure 2C**), which is 4 times the reported EC_50_ to activate PAR1 in HMEC.(12)

**Figure 2:**
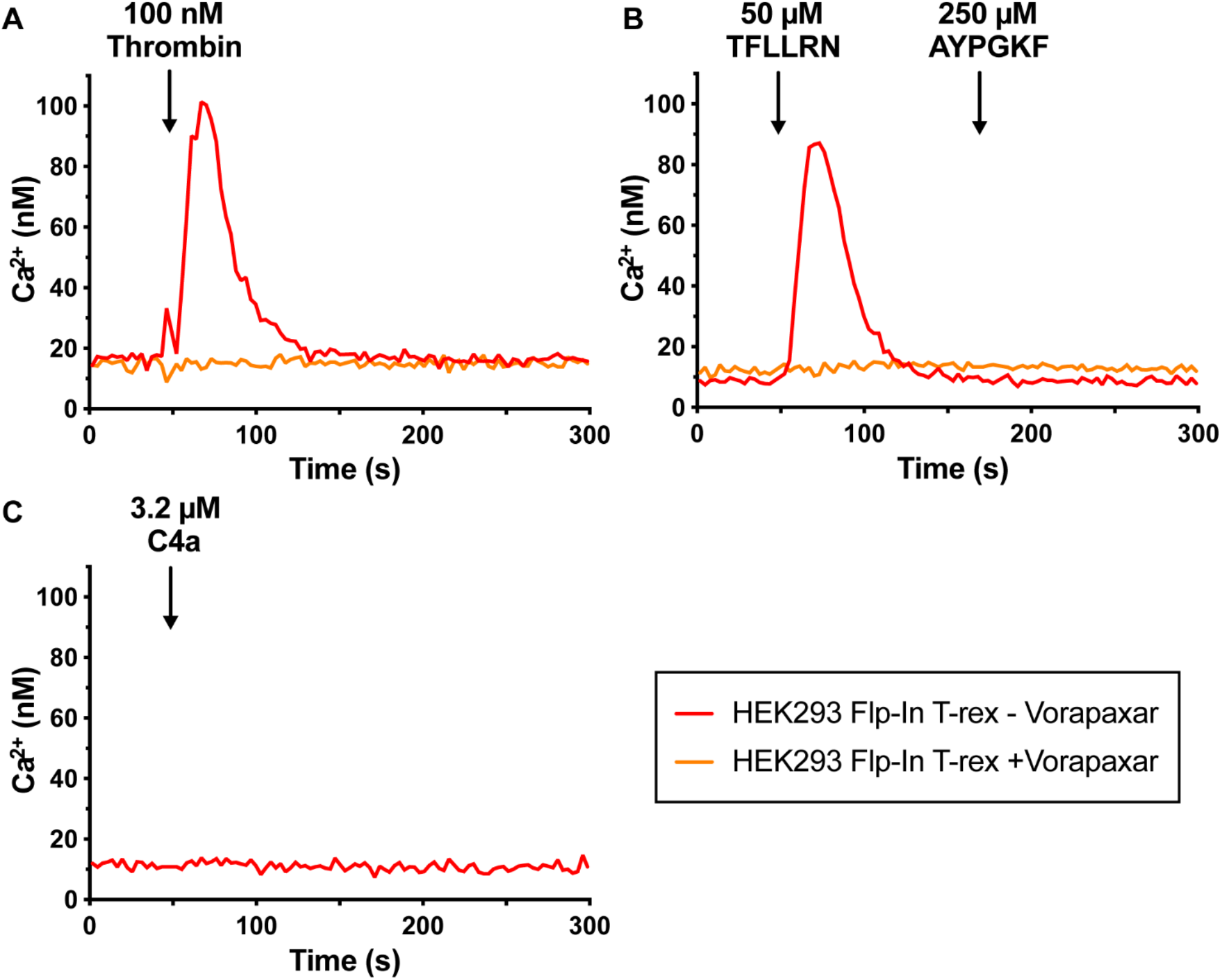
C4a did not induce PAR1 mediate intracellular calcium mobilization. **(A)** HEK293 cells express PAR1 endogenously, which respond to 100 nM thrombin stimulation (Red trace). The activation of PAR1 can be completely inhibited by pretreatment of 100 nM Vorapaxar (Orange trace). **(B)** The endogenous PAR1 on HEK293 can also be activated by 50 μM TFLLRN (PAR1-AP) (Red trace), which can be inhibited by vorapaxar (Orange trace). No endogenous PAR4 expresses on HEK293 and no intracellular calcium mobilization was triggered by 250 μM AYPGKF (PAR4-AP) (**C**) no PAR1-mediate calcium mobilization was triggered by 3.2 μM C4a.

To evaluate C4a-mediated PAR4 activation, we used inducible cell lines stably expressing wild-type PAR4 (PAR4-WT) with an N-terminal V5-epitope tag. To isolate the thrombin signaling to the exogenously expressed PAR4, cells were pretreated for 1 hr with 100 nM vorapaxar as above. PAR4 elicited the expected sustained Ca^2+^ mobilization compared to PAR1 stimulation (**Figure 3A**).(13, 17) Vorapaxar specifically ablated the response to PAR1, but left PAR4 response unaltered as determined by sequential activation by 50 μM of PAR1-AP stimulation (for endogenous PAR1) and 250 μM of PAR4-AP stimulation (for exogenous PAR4) (**Figure 3B**). As we observed in platelets, 2.5 μM C4a failed to induce Ca^2+^ flux in vorapaxar treated cells expressing PAR4 (**Figure 3C**).

**Figure 3:**
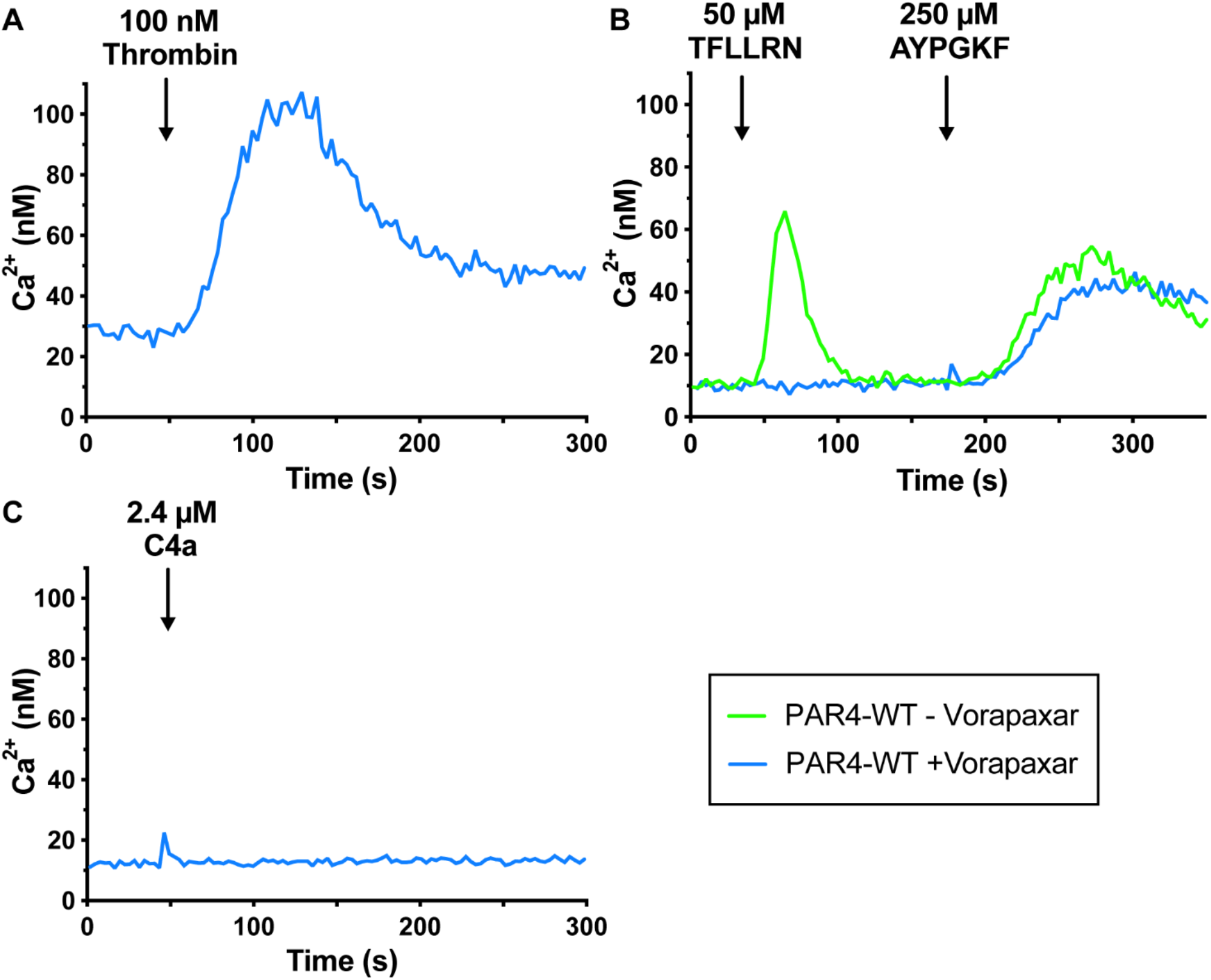
C4a did not induce PAR4 mediate intracellular calcium mobilization. **(A)** 100 nM Vorapaxar pretreated PAR4-WT cells were challenged with 100 nM thrombin and calcium mobilization responses to this stimulation was recorded. **(B)** The endogenous PAR1 on HEK293 can be activated by 50 μM TFLLRN (PAR1-AP) (Green trace), which can be inhibited by 100 nM Vorapaxar (Blue trace). The exogenously expressed PAR4 can be activated by 250 μM AYPGKF (PAR4-AP). (**C**) no PAR4-mediate calcium mobilization was triggered by 2.4 μM C4a.

The PAR family members share the same proteolytic activation mechanism; however, the sequence homology between family members is low and not significantly higher than other GPCRs. It is not surprising that agonist peptides derived from the tethered ligand and small molecule antagonist generally do not cross react between PARs. The notable exception is the cross reactivity of the PAR1 agonist peptide (SFLLRN) with PAR2. Interestingly, the PAR2 agonist peptide (SLIGKV) does not cross react with PAR1. Specific activation of PAR1 is achieved with the synthetic peptide TFLLRN. PAR1 and PAR4 are both activated by thrombin cleaving the N-terminus. However, the exodomains of PAR1 and PAR4 are dramatically different, which results in unique tethered ligands. The tethered ligand sequence for PAR1 is SFLLRNPNDKYE… compared to GYPGQVCANDSD… for PAR4. Since the canonical ligands for PAR1 and PAR4 are distinct and do not cross react, it implies that the ligand binding sites are also unique. Our recent hydrogen deuterium exchange data support this hypothesis.(13) Wang and colleagues show that C4a acts as a soluble ligand for both PAR1 and PAR4. These observations are exciting for two reasons. First, there have been no physiological soluble agonist for PAR1 or PAR4 previously described. Second, it strongly suggested that PAR1 and PAR4 have an additional ligand binding site that is conserved between PAR1 and PAR4 and regulates receptor function allosterically. Wang and colleagues screened C4a against a panel of GPCRs and identified C4a as a ligand for both PAR1 and PAR4 using a cell-based reporter assay. In human endothelial cells, their comprehensive experiments of ERK activation and calcium mobilization indicated that C4a-mediate cell activation is a PAR1- or PAR4-dependent manner, which suggested that C4a can act as a nontraditional free ligand that regulate both PAR1 and PAR4.(12)

We did not see activation platelets in response to C4a implying that it does not activate PAR1 or PAR4. C4a was also unable to active PAR1- or PAR4-mediated Gα_q_ activation on human embryonic kidney cell line (HEK293), which was measured by intracellular calcium mobilization. C4a concentration in plasma can be very different depending on the context, the disease condition, and sensitivity of the measurement methods. For example, the C4a concentration in the healthy control group is 68.91 ±33.23 ng/ml (7.97 nM) compared to the 108.48 ±83.83 ng/ml or 12.54 nM in the participants with neovascular age-related macular degeneration (nAMD) measured by a cytometric bead assay.(15) In another study using an ELISA, the concentration of C4a is 2398 ng/ml (277 nM) in the healthy controls compared to 2304 ng/ml (266 nM) in patients with chronic hepatitis B infection but without liver failure and 1811 ng/ml (209 nM) is the patient with hepatitis B infection related acute-on-chronic liver failure (ACLF).(16) However, in any case, treating human platelets with 3.2 μM of C4a should be more than enough to elicit platelet aggregation if C4a is an agonist for platelet PARs. The contradiction to the published results by Wang *et al.* further demonstrates the importance of cellular context for receptor activation. For example, PARs signal indirectly through Gα_i_ in platelets via feedback to P2Y12.(3) C4a may be allosterically activating PAR1 or PAR4 in microvascular cells to trigger Gα_i_ signaling. The mechanism by which cell selective activation of PARs by C4a is mediated is not known, but may involve specific membrane localization, cofactors, or both. Cell specific regulation of protease signaling by soluble mediators offers intriguing new lines of research to identify cofactors or membrane environments required. As the field moves forward, we can draw on the knowledge gained though investigating the APC-PAR1 axis which requires membrane localization and endothelial protein C receptor.

## Author Contribution

XH, MD and MN conceived the study, designed the experiments, performed the experiments and analyzed the data. XH and MN wrote the manuscript. MD critically read and edited the manuscript.

## References

1. Vu TK, Hung DT, Wheaton VI, Coughlin SR. Molecular cloning of a functional thrombin receptor reveals a novel proteolytic mechanism of receptor activation. Cell. 1991;64(6):1057–68.

2. Nieman MT. Protease-activated receptors in hemostasis. Blood. 2016;128(2):169–77.

3. Han X, Bouck EG, Zunica ER, Arachiche A, Nieman M. Protease-Activated Receptors. In: Michelson DA, editor. Platelets Fourth Edition. 4 ed: Academic Press; 2019. p. 243.

4. Zhao P, Metcalf M, Bunnett NW. Biased signaling of protease-activated receptors. Front Endocrinol (Lausanne). 2014;5:67.

5. Cottrell GS, Coelho AM, Bunnett NW. Protease-activated receptors: the role of cell-surface proteolysis in signalling. Essays Biochem. 2002;38:169–83.

6. Camerer E, Kataoka H, Kahn M, Lease K, Coughlin SR. Genetic evidence that protease-activated receptors mediate factor Xa signaling in endothelial cells. J Biol Chem. 2002;277(18):16081–7.

7. Sveshnikova AN, Balatskiy AV, Demianova AS, Shepelyuk TO, Shakhidzhanov SS, Balatskaya MN, et al. Systems biology insights into the meaning of the platelet’s dual-receptor thrombin signaling. J Thromb Haemost. 2016;14(10):2045–57.

8. Woulfe DS. Platelet G protein-coupled receptors in hemostasis and thrombosis. J Thromb Haemost. 2005;3(10):2193–200.

9. Voss B, McLaughlin JN, Holinstat M, Zent R, Hamm HE. PAR1, but not PAR4, activates human platelets through a Gi/o/phosphoinositide-3 kinase signaling axis. Mol Pharmacol. 2007;71(5):1399–406.

10. Peerschke EI, Yin W, Ghebrehiwet B. Platelet mediated complement activation. Adv Exp Med Biol. 2008;632:81–91.

11. Peerschke EI, Yin W, Ghebrehiwet B. Complement activation on platelets: implications for vascular inflammation and thrombosis. Mol Immunol. 2010;47(13):2170–5.

12. Wang H, Ricklin D, Lambris JD. Complement-activation fragment C4a mediates effector functions by binding as untethered agonist to protease-activated receptors 1 and 4. Proc Natl Acad Sci U S A. 2017;114(41):10948–53.

13. Han X, Hofmann L, de la Fuente M, Alexander NS, Palczewski K, INVENT, et al. PAR4 activation involves extracellular loop-3 and transmembrane residue Thr153. Blood. 2020;In Press.

14. Grynkiewicz G, Poenie M, Tsien RY. A new generation of Ca2+ indicators with greatly improved fluorescence properties. J Biol Chem. 1985;260(6):3440–50.

15. Lechner J, Chen M, Hogg RE, Toth L, Silvestri G, Chakravarthy U, et al. Higher plasma levels of complement C3a, C4a and C5a increase the risk of subretinal fibrosis in neovascular age-related macular degeneration: Complement activation in AMD. Immun Ageing. 2016;13:4.

16. Li Q, Lu Q, Zhu MQ, Huang C, Yu KK, Huang YX, et al. Lower level of complement component C3 and C3a in the plasma means poor outcome in the patients with hepatitis B virus related acute-on-chronic liver failure. BMC Gastroenterol. 2020;20(1):106.

17. Covic L, Gresser AL, Kuliopulos A. Biphasic kinetics of activation and signaling for PAR1 and PAR4 thrombin receptors in platelets. Biochemistry. 2000;39(18):5458–67.

